# A mechanism to prevent transformation of the Whi3 mnemon into a prion

**DOI:** 10.1101/2020.03.13.990119

**Authors:** Yasmin Lau, Iuliia Parfenova, Juha Saarikangas, Richard A. Nichols, Yves Barral, Fabrice Caudron

## Abstract

In response to deceptive courtship, budding yeast cells escape pheromone induced cell cycle arrest through coalescence of the G1/S inhibitor Whi3 into a dominant inactive super-assembly. Strikingly, Whi3 super-assemblies remain stable over many cell cycles in the mother cells and are not passed on to the daughter cells. Thereby, Whi3 coalescence encodes memory, conferring to it the property of a mnemon (Whi3^mnem^), a protein which conformational change maintain a trait that is permanent in the mother cell but is not inherited by daughter cells. Mnemons share structural features with prions, which are self-templating protein conformations that are inherited by daughter cells. Yet, how the maintenance and asymmetric inheritance of Whi3^mnem^ are achieved is unknown. Here, we report that Whi3^mnem^ is closely associated with endoplasmic reticulum (ER) membranes and retained in the mother cell by the presence of lateral membrane diffusion barriers at the bud neck. Strikingly, barrier defects made Whi3^mnem^ propagate in a mitotically stable manner, like a prion. Alike Whi3^mnem^, transformation of Whi3 into a prion required its poly-glutamine prion-like domain. Thus, we propose that Whi3^mnem^ is in a self-templating state, lending temporal stability to the memory that it encodes, while its anchorage into the compartmentalized membranes of the ER ensures its confinement in the mother cell and prevents its infectious propagation. These results suggest that confined self-templating super-assembly is a powerful mechanism for the long-term encoding of information.

## Introduction

Prions are proteins that can adopt several conformations with at least one of them that is self-templating, lending it a self-perpetuating character. Prion conversion is essential for physiological processes including innate immunity (Hou et al., 2011) and the generation of phenotypic diversity of single celled organisms ranging from bacteria to yeast (Coustou et al., 1997; True and Lindquist, 2000; Yuan and Hochschild, 2017). Beyond physiology, prions were initially discovered as agents of devastating diseases in humans (Kuru, Creutzfeldt-Jakob’s disease) and animals (Scrapie and Bovine spongiform encephalopathy) (Prusiner, 1982). Moreover, prion-like behavior of different proteins may be linked to other diseases including Alzheimer’s and Parkinson’s diseases (Goedert, 2015), as well as resistance to anti-fungal drugs (Suzuki et al., 2012). What mechanisms control prions and particularly their self-templating activity is still an open question.

Prions are common protein elements and several have been identified in the budding yeast *S. cerevisiae* (Wickner et al., 2015) allowing a wealth of research to understand their regulation and function (Harvey et al., 2018). The protein Sup35 can adopt a conformation to form the [*PSI*+] prion, the best characterized to date (Tuite and Cox, 2006). Sup35 is a translation terminator that loses its function in its [*PSI*+] form, resulting in stop codon readthrough and phenotypic diversity (True and Lindquist, 2000). Importantly, Sup35 contains a prion-like domain, defined as rich in Glutamine and Asparagine (Q/N) residues and depleted of charged and hydrophobic residues (Alberti et al., 2009). This domain drives the Sup35 prion state by adopting a self-templating conformation (Serio et al., 2000) and mediating the assembly of Sup35 into amyloid fibrils. Once established, fibrils formed by the Sup35 protein are fragmented into smaller seeds by the protein disaggregase Hsp104 (Narayanan et al., 2006). The self-templating activity of prion seeds converts most of the protein pool to the prion form and the seeds are free to diffuse to the daughter cells. Therefore, prions behave as infectious particles, establishing [*PSI*+] not only in the cell in which it originated but also in all its daughter cells and hence the growing colony. In silico analysis of the yeast proteome has identified around 200 proteins in *S. cerevisiae* containing a potential prion-like domain (Alberti et al., 2009). However, we still do not know if all prion-like domains can adopt self-templating conformations and whether all these proteins function as prions.

Whi3 contains two prion-like domains, composed of PolyQ- and PolyN-rich streches (Alberti et al., 2009; Caudron and Barral, 2013). Whi3 is an mRNA binding protein involved in the timing of the G1/S transition of the cell cycle and the control of cell size in *S. cerevisiae* (Garì et al., 2001). The function of Whi3 that has been studied the most is the regulation of the G1/S transition of the cell cycle through binding and inhibiting the translation of the mRNA that encodes the G1 cyclin *CLN3*. This inhibition ensures that entry into S phase is delayed until cells reach a critical size. The prion-like domains of Whi3 are required for Whi3 to adopt a conformation that releases Whi3’s inhibition on *CLN3* mRNA and allows a long-lasting phenotypic switch. Upon exposure to mating pheromone, haploid yeast cells arrest in the G1 phase of the cell cycle and grow towards the source of pheromone. This cytoplasmic projection is termed a shmoo. After prolonged exposure of pheromone without a mating partner in reach, yeast cells become refractory to the pheromone signal and resume their cell cycle (Caudron and Barral, 2013; Moore, 1984). They switch from a shmooing phase to a budding phase. Remarkably, once established, this pheromone refractory state is stable, lending memory to the cell that there is no partner available. As a consequence, these cells keep on budding for the remainder of their life span even in the presence of mating pheromone. Strikingly, daughter cells do not inherit this adaptation and restore their ability to shmoo in response to pheromone upon separation from their mother cell (Caudron and Barral, 2013). Whi3’s prion-like domains trigger both escape from pheromone arrest and the memory state by mediating the condensation of Whi3 into super-assemblies.

Whi3s’ domain organization and ability to switch between soluble and condensed phases are features common to prions such as Sup35. The pheromone refractory state is very stable once it has been established, even when pheromone is removed from the medium. The deletion of either prion-like domain of Whi3 results in a late escape from pheromone arrest that is not stably maintained in the mother cells. In other words, cells can switch back and forth to a shmooing and a budding phase in the presence of pheromone. Moreover, when cell extracts are run on a semi-denaturing gel, Whi3 appears to be mostly monomeric in untreated cells and mostly found in large, homo-multimeric assemblies in cell treated with pheromone (Caudron and Barral, 2013). These data point to a possible self-templating activity of Whi3 in cells that are exposed to pheromone.

Importantly, despite many shared features with known prions, the mode of inheritance of Whi3 super-assemblies is different from the inheritance of prions during cell division (Caudron and Barral, 2013; Tuite, 2016). Indeed, the Whi3 super-assemblies remain in the mother cells at mitosis, ensuring that the memory state is stable in the mother cell and is not propagated to the progeny. On the contrary, prions infect the bud of a dividing yeast cell. Given its role in cellular memory and its behavior being so distinct from prions, Whi3 was termed a mnemon (Whi3^mnem^).

While the prion-like domains of Whi3^mnem^ are important for escape from pheromone arrest, we do not know the mechanism of memory stability and if this involves a self-templating activity of Whi3^mnem^. If this is the case, what makes mnemons distinct from prions is unclear. In this work, we wondered whether the confinement of Whi3^mnem^ to the mother cell requires the presence of lateral membrane diffusion barriers. Diffusion barriers are membrane specialized domains that limit the diffusion of membrane associated structures across cellular appendages (Caudron and Barral, 2009; Saarikangas and Barral, 2011) including the primary cilium (Hu et al., 2010; Kim et al., 2010), dendritic spines (Ewers et al., 2014) and the sperm tail (Toure et al., 2011). In budding yeast, diffusion barriers form at the bud neck in the ER membranes and the nuclear envelope during closed mitosis. Their formation depends on septins, a family of cytoskeletal protein forming filaments (Mostowy and Cossart, 2012) at the bud neck (Luedeke et al., 2005), proteins involved in sphingolipid biosynthesis including the sphinganine C4-hydroxylase Sur2 (Clay et al., 2014) and proteins involved in polarized cell growth such as the actin nucleation promoting factor Bud6 (Graziano et al., 2013, 2011) and the small GTPase Bud1/Rsr1 (Bender and Pringle, 1989). Diffusion barriers in yeast limit the diffusion of ageing factors, such as nuclear pores, misfolded proteins, age-induced protein deposits and DNA circles from the mother cell to the bud (Clay et al., 2014; Denoth-Lippuner et al., 2014; Saarikangas et al., 2017; Shcheprova et al., 2008).

In light of these data, we first asked whether membrane diffusion barriers contribute to Whi3^mnem^ asymmetric behaviour during cell division.

## Results

### Cells lacking diffusion barriers can acquire a novel pheromone refractory state

Growth of yeast colonies was monitored on solid medium containing low concentration of pheromone (10nM). Wild type cells grew slowly while cells that hardly escape pheromone arrest (*whi3*-Δ*pQ*, (Caudron and Barral, 2013)) grew very poorly (Figure 1A). Both strains grew equally well on a solid medium that does not contain pheromone. On medium containing pheromone, wild type cells initially shmoo and resume cell division after several hours. Their daughter cells behave similarly, first shmooing and then resuming cell division. This slows down the production of daughter cells and explains why wild type cells grow slowly in the presence of pheromone. In contrast, *whi3*-Δ*pQ* mutant cells keep on shmooing for a very long time before resuming cell division and many cells will not even escape pheromone induced cell cycle arrest. Therefore, *whi3*-Δ*pQ* mutant cells grow even slower on pheromone containing plates. To test for a role of diffusion barriers in the compartmentalization of the pheromone refractory state, we monitored the growth of several mutant strains with impaired lateral diffusion barriers. We hypothesized that if diffusion barriers are involved in the confinement of Whi3^mnem^ these mutant strains may grow differently on pheromone. Some of the *bud1*Δ, *bud6*Δ and *sur2*Δ mutants grew as wild type and other much better on pheromone containing plates, even at high pheromone concentration (0.6μM, Figure 1A). At such a high concentration of pheromone, cells cannot escape pheromone arrest, suggesting that colonies growing in such conditions had acquired a strong resistance to pheromone. Colonies growing on high pheromone concentration kept their resistance to pheromone after several rounds of streaking; we termed these isolates as constitutive escapers (CE). To measure how frequent CE are, we plated at least 44 independent clones for each mutation obtained from heterozygous diploid on rich media that do not contain pheromone or contain a high pheromone concentration (0.6μM) (Figure 1A-B). Frequency of CE appearance was highly variable between individual clones of wild type strains (between 1.11 × 10^−7^ and 5.3 × 10^−2^) with a median of 1.36 × 10^−5^ cells. Phenotypic mutation rate for sterile mutants was measured in a *bar1*Δ background at 3.07 × 10^−6^/genome/generation (Lang and Murray, 2008). We found that it was also variable in *sur2*Δ clones but the median was significantly increased 3.13 times compared to wild type (4.26 × 10^−5^, Figure 1B). In *bud6*Δ and *bud1*Δ mutant cells, frequencies again varied extensively between clones and the median frequency was significantly increased 6.07 times (8.27 × 10^−5^) and 3.51 times (4.78 × 10^−5^) respectively, compared to wild type. We analyzed these results differently in an attempt to standardize the variances. The ratios were logit transformed (see material and methods for more details). Having fitted the average for each yeast strain, the residuals were clearly strongly asymmetrical (Supplemental Figure 1). This pattern might be expected if in a subset of cases the constitutive escaper phenotype occurred early in the culture. Since the incidence of these outcomes did not differ between strains, they were excluded from subsequent analysis of the ratios. Figure 1C shows the distribution of the logit transformed ratios for each genotype. A linear model describing the means of each strain, showed a highly significant difference between wild type and diffusion barrier mutants (ANOVA p<2e-16). These data suggest that the frequency increase of constitutive escapers is related to the disruption of the diffusion barrier. Moreover, the fact that we measured very variable frequencies and many of them being well above mutation rate suggest that CE are induced by an epigenetic mechanism. Since escape from pheromone arrest involves Whi3^mnem^, we considered the possibility of a prion being the molecular basis of the CE phenotype.

**Figure 1.**
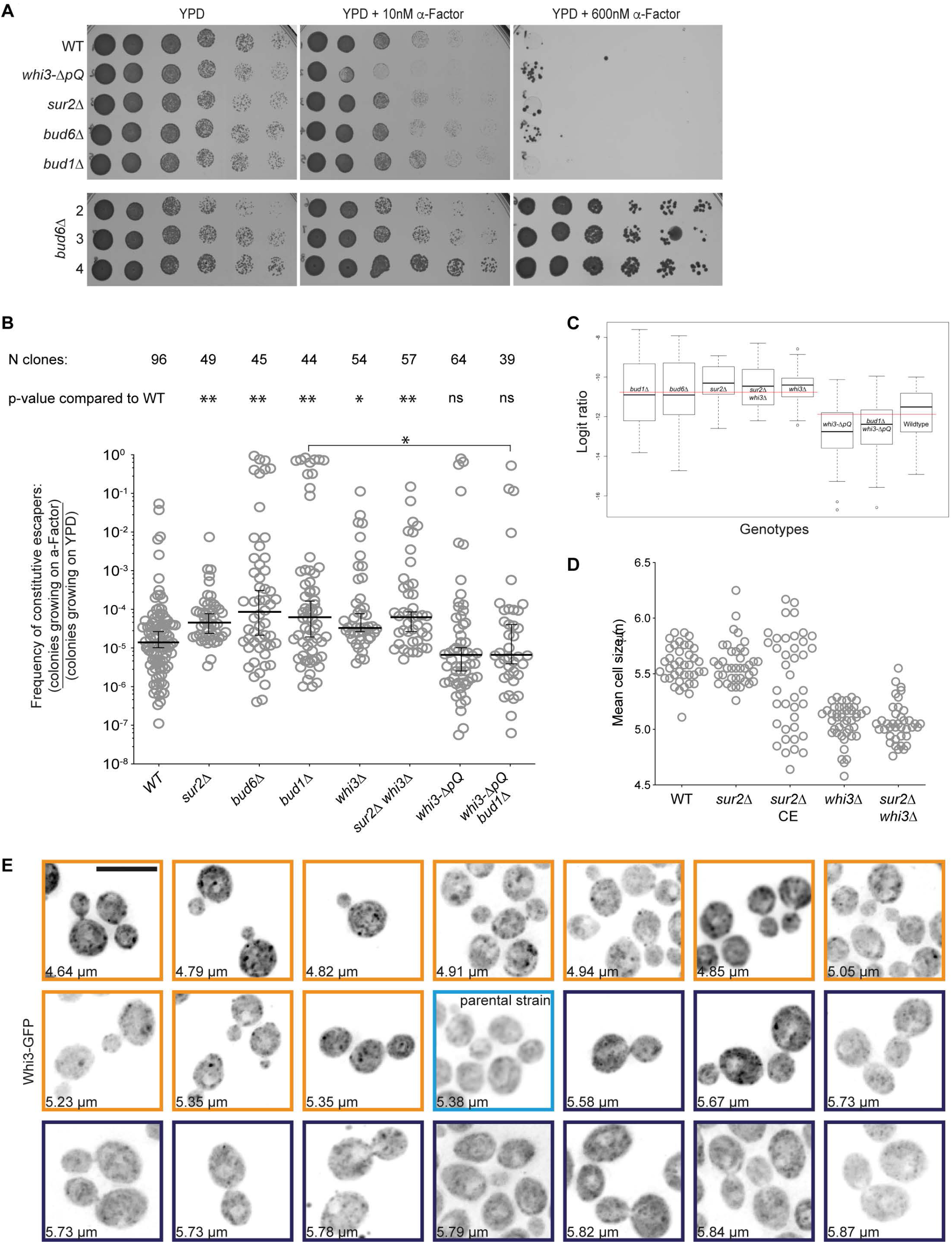
A novel epigenetic phenotype, constitutive escapers. (A) Serial 1/10 dilutions of exponentially growing cultures of indicated strains spotted on a YPD solid medium (left) or YPD containing α-factor (10nM middle and 600nM right). The bottom panel shows three *bud6*Δ independent clones with high frequency of CE. (B) Frequency of appearance of CEs in the indicated genotypes (median with 95% confidence interval, n>37 clones for each, Dunn’s multiple comparisons test was used to determine significance). (C) Distribution of the logit transformed ratios (R) for each genotype. (D) Cell size distributions of individual clones or CE isolates of indicated strains (n>39 clones for each). (E) Maximal projection images of *sur2*Δ cells expressing Whi3-GFP from 20 CE isolates and the parental strain. Isolates are in order of mean cell size from the smallest (orange frame are smaller than the blue framed parental strain) to the largest (purple are larger than the parental strain). Scale bar = 5μm.

Upon microscopic observation of CE, we noticed that some CE isolates seemed to have a small cell size. Therefore, we measured the cell size of several CE. While wild type cells and *sur2*Δ parental strains had comparable cell size (5.6μm, Figure 1D), *sur2*Δ^*CE*^ displayed isolates which were on the large range or on the small range of cell sizes. Remarkably, the small cell sized *sur2*Δ^*CE*^ were comparable in size to cells deleted for *WHI3* either in a wild type or in a *sur2*Δ background (Figure 1D). This prompted us to ask whether Whi3 could be a prion in CE isolates and if it was required for the formation of CE. Classically, prion forms of proteins lose the function of the native conformation. Thus, if Whi3 was losing its function in a prion form, we would expect that deleting *WHI3* would result in a CE phenotype. The frequency of CE in *whi3*Δ mutant strains was increased compared to wild type (2.45 times, 3.33 × 10^−5^) and further increased in *whi3Δ sur2*Δ strains (4.58 times, 6.23 × 10^−5^). This result indicate that Whi3 inactivation is not sufficient to make cells refractory to pheromone, although it may contribute to it. We next wondered whether Whi3 was required to maintain the CE phenotype. We supposed that if Whi3 was acting as a dominant negative form in these cells, deleting *WHI3* in *sur2*Δ^*CE*^ would restore pheromone arrest. We specifically deleted *WHI3* in several *sur2*Δ^*CE*^ that had a small cell size. This deletion did not restore pheromone arrest (not shown) suggesting that maintenance of the CE phenotype is not dependent on Whi3 or that Whi3 super-assemblies are not the only dominant factor repressing pheromone response in CEs. Altogether, we observe that *whi3*Δ cells are not all CE (most of the cells arrest upon exposure to pheromone) yet deleting *WHI3* seems to favor CE formation. However, Whi3 is not required to maintain the CE phenotype. One interpretation of these results may be that *WHI3* inactivation is one step for cells to become CE and that somehow, the diffusion barrier inhibits the transition to the CE state.

To further test whether Whi3 may be in a prion form in CE, we isolated 20 CE from a *sur2*Δ parental strain (*sur2*Δ^*CE*^) expressing Whi3-GFP from its endogenous locus on plates containing high pheromone concentration and we observed the localization of Whi3-GFP in these cells. In parallel, we obtained cell size measurements of these isolates. In the parental *sur2*Δ strain, Whi3-GFP localized rather diffusely throughout the cytoplasm and to a few granules ((Caudron and Barral, 2013; Garì et al., 2001) and Figure 1E). In small cell sized *sur2*Δ^*CE*^, Whi3 localized diffusely throughout the cytoplasm and to brighter foci that were substantially bigger and more intense than the granules shown by the parental strain. These bright foci were much less frequent in large cell sized CE (Figure 1E). Remarkably, these bright foci localized both to the mother and bud compartments of dividing cells. This is in contrast to Whi3^mnem^ super-assemblies which are formed in the mother cell and do not typically appear in the buds (Caudron and Barral, 2013). Therefore, in small cell sized *sur2*Δ^*CE*^, Whi3 tends to adopt a localization pattern reminiscent to that of the classical yeast prion Sup35, in its [*PSI*+] form (Derdowski et al., 2010). We contemplated the possibility that if Whi3 needs to be in a prion form for cells to become CE, it would require its prion-like domain. To test this hypothesis, we estimated the frequency of CE in *whi3*-Δ*pQ* mutant clones. CE frequency was remarkably lower in these cells than in any other strain we tested (6.25 × 10^−6^, 2.18 times smaller than wild type strains, Figure 1B, and significantly lower than wild type using the logit transformed data, p<0.0008, Figure 1C). Deleting *BUD1* in a *whi3*-Δ*pQ* strain did not increase the frequency of CE much (7.45 × 10^−6^, 1.192 times higher than in *BUD1 whi3-ΔpQ* mutant strain) and was lower than in *bud1*Δ mutant cells.

Altogether, we found that disruption of the diffusion barrier increases the appearance of CE, that many of them are small cell sized and display a localisation of Whi3 to bright foci. The formation of CE is increased when *WHI3* is deleted but decreased in strains that lack the prion-like domain of Whi3. This suggest that Whi3 is indeed in a prion form and that inactivation of Whi3 is a major contribution to the formation of CE. However, since the frequency of CE can be still very high in CE that lack the prion-like domain of Whi3, it is very likely that other factors are involved, possibly novel, yet undiscovered, prions.

To test whether the molecular events leading to the formation of CE is indeed caused by prions, we tested how growth on high pheromone concentration segregated at meiosis. Prions segregate in a non-Mendelian manner at meiosis and are usually passed on to all of the four meiotic products. Consistent with their extensive resistance to pheromone treatment, crossing the CE to wild type cells was inefficient. However, we could backcross some of them and dissect the obtained tetrads. Meiosis gives rise to 4 spores, 2 of which are *MATα* and 2 *MAT***a**. Therefore, we expected that out of 4 spores, 2 would always grow on alpha-factor because they are *MATα*. In wild type cells, the 2 other spores do not grow on alpha-factor (0.6μM, Supplemental Figure 2). If in the CE more than 2 spores were growing, it would mean that they had inherited the CE phenotype. If the CE is due a mutation that is unlinked to the *MAT* locus, 4/6 of the tetrads should have 3 spores growing on alpha-factor containing medium, 1/6 should contain 4 such spores and the last 1/6 of the tetrads should contain only 2 of them (pattern #1). If it is due to a mutation linked to the *MAT* locus, the fraction of tetrads with 3 and 4 spores growing on alpha-factor containing medium should be increased (pattern #2). In case of a non-mendelian factor propagating through meiosis, all tetrads should contain 4 spores growing on alpha-factor containing medium (pattern #3). Finally, a non-mendelian factor that does not pass meiosis should produce tetrads with always only 2 spores growing on alpha-factor containing medium (pattern #4). We backcrossed 13 independent CE strains and tested the growth of each spore on alpha-factor (0.6μM). For 4 backcrosses, we observed tetrads in which 2, 3 or 4 spores out of 4 were growing on α-factor (Supplemental Figure 2). These backcrosses fall in the pattern #1, suggesting the presence of a single ‘sterile’ mutation segregating independently from the mating type locus. In 4 other backcrosses, the majority of the tetrads contained 2 spores and few 3 spores growing on alpha-factor containing medium, most compatible with the pattern #4. Furthermore, 5 backcrosses followed strictly pattern #4 (Supplemental Figure 2). Thus, the last 9 backcrosses, which are not compatible with a single mutation, suggests that the CE phenotype is due to non-mendelian factor that is lost during meiosis. To further test if these CE traits are based on a prion-like mechanism, we tested whether they were cured upon inhibiting different prion effectors. We isolated 31 *sur2*Δ^*CE*^ and passaged them three times on YPD, YPD supplemented with guanidine hydrochloride (3 mM) to inhibit Hsp104 (Ferreira et al., 2001; Tuite et al., 1981) or YPD supplemented with radicicol (10μM) to inhibit Hsp90 (Sharma et al., 1998). In addition, we transformed all 31 *sur2*Δ^*CE*^ and the parental *sur2*Δ strain with a dominant negative allele of *SSA1* (*SSA1*^*DN*^, (Brown and Lindquist, 2009; Jarosz et al., 2014)). In all cases, after 3 passages the 31 *sur2*Δ^*CE*^ were still able to grow on YPD containing pheromone (0.6μM), while the parental *sur2*Δ strain was not (Supplemental Figure 3). However, upon microscopic observation of *sur2*Δ^*CE1*^, we found that many cells were shmooing and other dividing. This was not the case for other *sur2*Δ^*CE*^, and it was also not the case for *sur2*Δ^*CE1*^ passaged on YPD without drugs or with GuHCl or radicicol (Supplemental Figure 3). Therefore, the CE phenotype is not cured by either GuHCl, radicicol or passages for many generations on YPD, but one variant was partially cured upon inhibition of the Hsp70 chaperone Ssa1.

**Figure 2.**
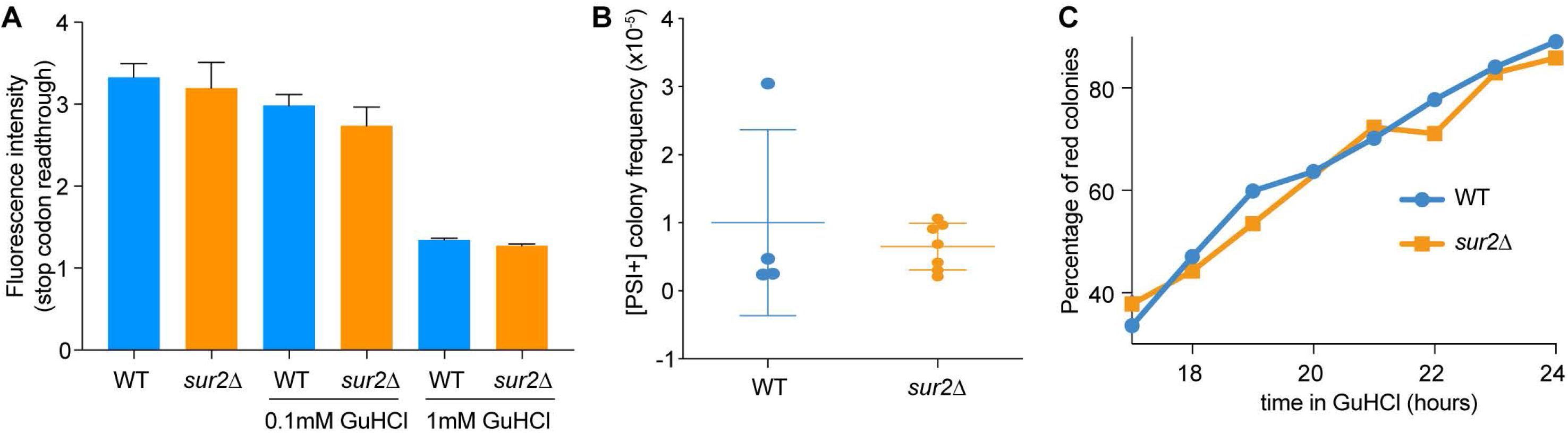
Endoplasmic reticulum compartmentalization by a lateral membrane diffusion barrier is not required for prion induction and curing. (A) Fluorescence intensity measured by flow cytometer of [PSI+] WT and [PSI+] *sur2*Δ cells treated or not with 0.1mM and 1mM GuHCl. (B) Frequency of *de novo* [PSI+] appearance and (C) percentage of cells cured of [PSI+] over time. Graphs A and B display mean ± SD.

**Figure 3.**
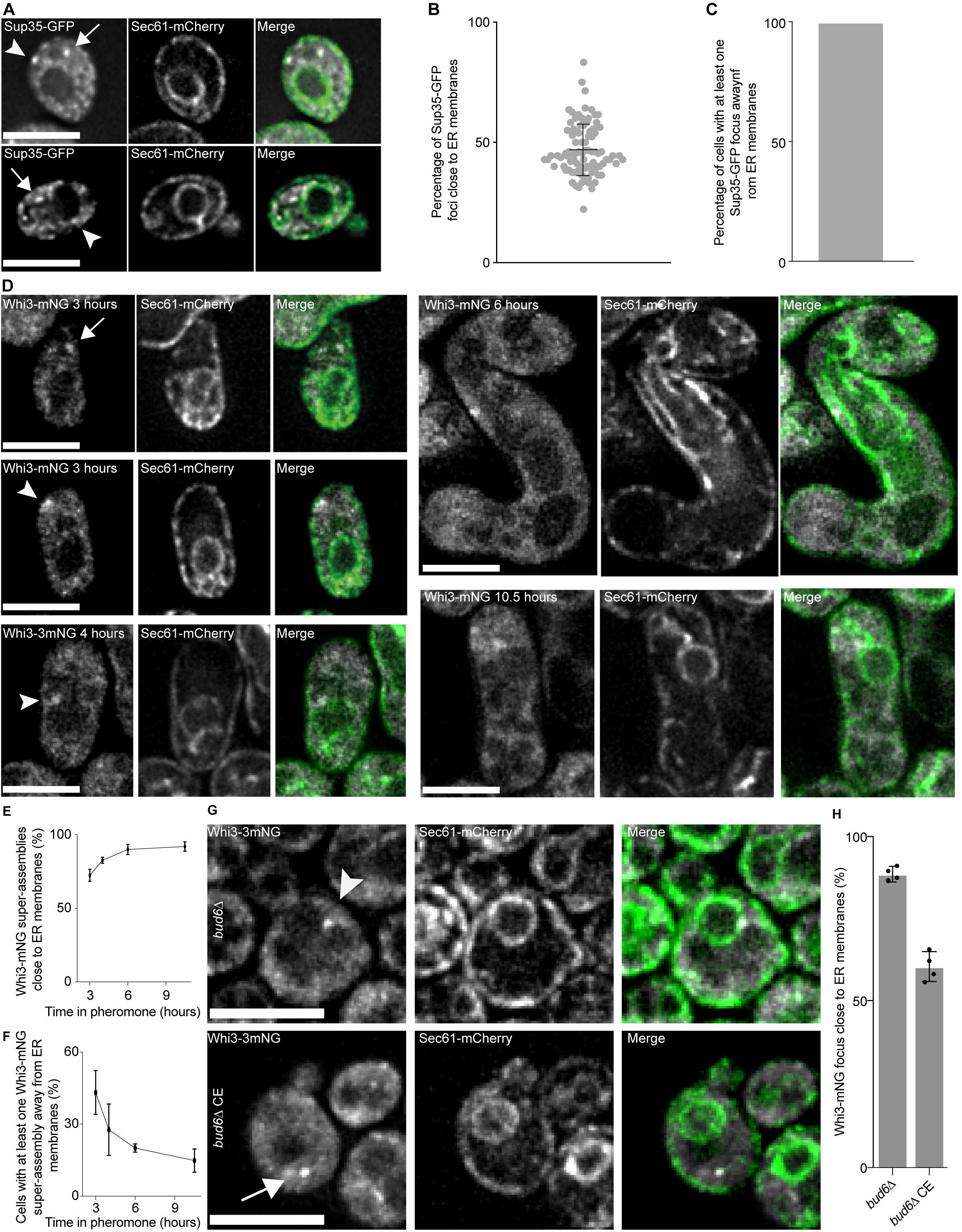
Sup35 and Whi3 prion foci are not closely linked to ER membranes while Whi3 super-assemblies and granules are. (A) Single focal plane images of [*PSI*+] cells expressing Sup35-GFP and Sec61-mCherry. (B) Percentage of Sup35 foci close to ER membranes (1180 foci analysed from 90 cells of three independent clones, mean ±SD is presented). (C) Percentage of cells with at least one Sup35-GFP focus away from ER membranes (3 clones with more than 90 cells each). (D) Single focal plane images of cells expressing Whi3-mNG and Sec61-mCherry exposed to pheromone for 3 hours (top left and middle left) and 4 hours (bottom left), and cells exposed for 5 hours to pheromone and released in a pheromone free medium then imaged at 6 hours (top right) and 10.5 hours (bottom right) after initial exposure to pheromone. (E) Percentage of Whi3-mNG super-assemblies close to ER membranes (3 independent clones, >300 super-assemblies from >153 cells were analysed, mean ±SD are presented). (F) Percentage of cells with at least one Whi3-mNG super-assembly away from ER membranes (3 independent clones, >300 super-assemblies from >153 cells were analysed, mean ±SD are presented). (G) Single focal plane images of *bud6*Δ cells (top) and *bud6*Δ CE cells (bottom) expressing Whi3-mNG and Sec61-mCherry. (H) Percentage of Whi3-mNG granules (*bud6*Δ) and foci (*bud6*Δ CE) localizing close to ER membranes (4 clones or isolates, 527 granules and 359 foci in 200 cells were analysed, mean ±SD are presented). For all panel, arrows point at foci away from the ER and arrowheads point at foci close to the ER. Scale bars = 5 μm.

Altogether, our data suggest that the majority of the CE formed in cells lacking a diffusion barrier at the bud neck are due to a non-mendelian factor and that for about at least half of them Whi3 is contributing to their formation by behaving not anymore as a mnemon but as a prion.

### Endoplasmic reticulum compartmentalization is not required for prion induction and curing

Because diffusion barriers seem to play a role in the transformation of Whi3 into a prion, we next asked whether diffusion barriers also play a role in prion induction and curing. We specifically focused on the best studied prion in yeast, [*PSI*+]. We previously observed that farnesylation of the Hsp40 co-chaperone Ydj1 was required to lower premature stop codon read-through in the sequence coding for GFP in [*PSI*+] cells (Saarikangas et al., 2017). Strength of codon read-through was linked to the potential collection of prion seeds to ER membranes. We therefore analyzed stop codon readthrough in *sur2*Δ cells. Using flow cytometry, the fluorescence intensity of strains that express a GFP allele containing a premature stop codon was measured (as in Saarikangas et al., 2017). Deletion of *SUR2* did not change the fluorescence intensity of these cells (Figure 2A). We next probed for a role of lateral diffusion barriers in the de novo appearance and curing of [*PSI*+] prions. There was no significant difference (p=0.5204, t-test) in the [*PSI*+] appearance frequencies in wild type and *sur2*Δ strains (Figure 2B). We also analyzed the dynamics of [*PSI*+] curing by Guanidine Hydrochloride (GuHCl, 3 mM) in wild type and *sur2*Δ cells and observed very similar dynamics of curing (Figure 2C). Moreover, stop codon readthrough was comparably lower in wild type and *sur2*Δ cells treated with 0.1mM GuHCl and 1mM GuHCl demonstrating that we could quantitatively measure the strength of stop codon readthrough in these experiments (Figure 2A). Altogether, these data establish that compartmentalization of endo-membranes by a diffusion barrier at the bud neck does not control the inheritance of Sup35 prion seeds during cell division.

### Whi3 super-assemblies and Sup35 foci differ in their association to ER membranes

Because diffusion barriers affect Whi3 transformation into a prion and not [*PSI*+], we wondered if Sup35 and Whi3 associated to different extends with ER membranes. We first analyzed how Sup35-GFP, in its [*PSI*+] prion form, localized relative to the ER visualized using the translocon subunit Sec61 tagged with mCherry as a marker. Sup35-GFP formed several small foci in dividing cells and we found that most of these foci were spatially excluded from the Sec61 signal (Figure 3A and B, 53.1 ±10.7% of Sup35-GFP foci were away from the Sec61-mCherry signal). We also noticed that 100% of the cells contained at least one focus away from ER membranes (Figure 3C). This may explain why curing and induction of [*PSI*+] is not affected in *sur2*Δ mutant cells. Next, we analyzed the localization of Whi3-mNeonGreen (Whi3-mNG) in its mnemon form relative to Sec61-mCherry. Contrarily to Sup35-GFP, in cells that were exposed to pheromone for 3 hours Whi3-mNG super-assemblies were mostly apposed to ER membranes (72.5 ±4.1% of super-assemblies, Figure 3D and E). Moreover, the fraction apposed to the ER system increased upon longer times of pheromone exposure (82.7 ±2.1% after 4 hours, 90.0 ±3.5% after 6 hours and 92.1 ±3.2% after 10.5 hours, Figure 3D and E). Remarkably, while 41.2 ±9.1% of the cells had at least one Whi3-3GFP super-assembly away from the ER after 3 hours of pheromone treatment, this value dropped to 27.7 ±10.7%, 20.0 ±1.7% and 14.8 ±4.9% after 4 hours, 6 hours and 10.5 hours of pheromone treatment respectively (Figure 3F). Finally, we analyzed the localization of Whi3-mNG relative to ER membranes in small *bud6*Δ^*CE*^. We found that 60.27% of the Whi3-mNG foci were close to ER membranes in *bud6*Δ^*CE*^ cells, in stark contrast to the granules in non-CE *bud6*Δ cells which were much more strongly associated to ER membranes (88.17%, Figure 3G and H).

Altogether, we conclude that association of Whi3^mnem^ with ER membranes is tighter with time after escape form pheromone arrest and that a major difference between Whi3^mnem^ super-assemblies and the prion form of Sup35 or the Whi3 foci in CE is their different link to the ER membranes network.

### Endoplasmic reticulum compartmentalization is required for the retention of Whi3^mnem^ and the pheromone refractory state in the mother cell

The establishment and the maintenance of the pheromone refractory state is facilitated by Whi3’s conformational change and super-assembly formation. We wondered whether mother cells with defective diffusion barriers were more likely to pass Whi3 super-assemblies to their daughters. Therefore, we treated cells expressing *WHI3* fused to 3 superfolding Green Fluorescent Proteins in tandem (Whi3-3GFP) with pheromone for 5 hours and then released them in a pheromone free medium for 1.5 hours to allow mother cells to produce a bud. The mother and bud pairs were imaged and the localization of Whi3-3GFP analyzed. Most wild type mother cells contained a super-assembly of Whi3^mnem,^ and we observed similar results in *bud6Δ, sur2*Δ and *bud1*Δ mutant cells. However, while only 24.00 ±6.22% of the wild type buds had a super-assembly, 62.24 ±3.85% of the *bud6*Δ, 64.01 ±3.65% of the *sur2*Δ and 65.26 ±4.15% of the *bud1*Δ mutant buds already had at least one (Figure 4A-B). These data are consistent with a role for diffusion barriers in the retention of Whi3^mnem^ super-assemblies in the mother cells that escaped pheromone arrest.

**Figure 4.**
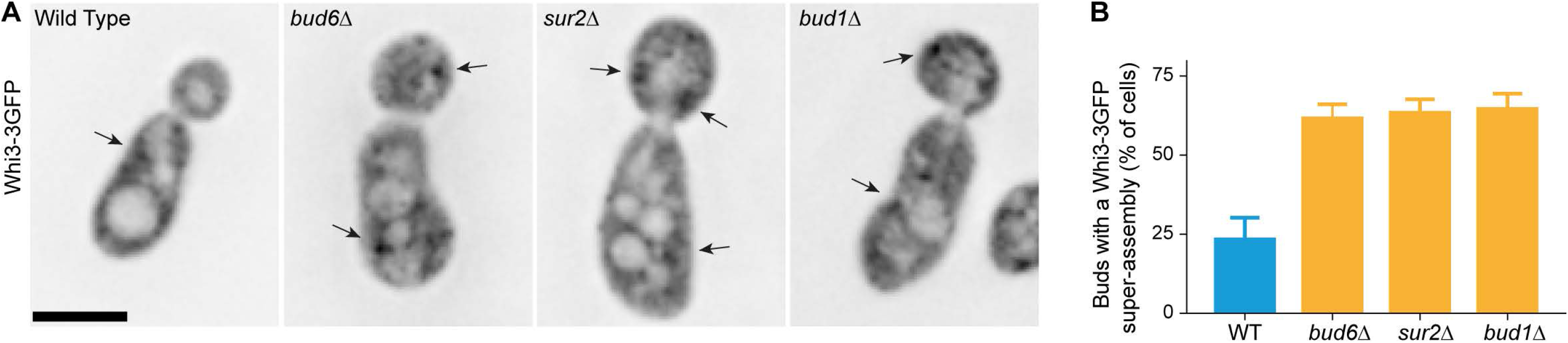
Whi3-3GFP super-assemblies form in the buds of mutants with a weak diffusion barrier. (A) Single focal plane images of Whi3-3GFP expressing cells. Scale bar = 5μm. (B) Quantification of buds with a detectable Whi3-3GFP super-assembly (n=122 cells, 148 cells, 180 cells and 165 cells for wild-type, *bud6*Δ, *sur2*Δ and *bud1*Δ cells respectively) Results from *bud6Δ, sur2*Δ and *bud1*Δ are significantly different from wild type (p<0.0001, one-way ANOVA).

We then tested whether diffusion barriers are required for the retention of the pheromone refractory state in the mother cell. We exposed haploid *MAT***a**cells to pheromone as before (Caudron and Barral, 2013). Wild type cells initially all shmooed upon 7 nM α-factor treatment and 90.5% of the cells escaped pheromone arrest with an average timing of 7.35 ± 3.00 hours (Figure 5A-B). Mother cells kept this refractory state faithfully, 100% of the initial mother cells did not produce a second shmoo after escaping pheromone arrest. We previously found that roughly 90% of the daughter cells from mothers that have escaped from pheromone arrest undergo shmooing in response to pheromone, upon separation from their mother in the presence of pheromone (Caudron and Barral, 2013). We refined our analysis here by separating data depending on whether daughters were the first or subsequent daughters after escape from pheromone arrest. We observed that nearly 50% of the first daughter cells fail to respond to pheromone after birth, while this number drops to 14.2%,13.3%, 6.9% and 4.4% for the subsequent 2^nd^, 3^rd^, 4^th^ and 5^th^ daughters of the same mother cell (Figure 5C). In order to assess whether the retention of the pheromone refractory state in the mother cell is indeed dependent on the presence of diffusion barriers at the bud neck, we tested *sur2*Δ, *bud6*Δ and *bud1*Δ mutant cells. For reasons that remain unknown at this stage, cells with a *sur2*Δ mutation escaped from pheromone arrest slightly earlier than wild type cells (5.9 ± 2.0 hours, Figure 5 A-B). Once these mother cells escaped, they maintained the pheromone refractory state as efficiently as wild type cells and did not shmoo again. However, the fraction of *sur2*Δ daughter cells that failed to shmoo after separating from their mother cell was increased 1.51, 2.83 and 1.66 folds for the 1^st^, 2^nd^ and 3^rd^ daughters respectively. (Figure 5C). The *bud6*Δ mutant cells responded to pheromone as well as wild type cells and escaped pheromone arrest slightly later than them (Figure 5A-B). As for the *sur2*Δ daughter cells, many *bud6*Δ daughter cells failed to shmoo upon separation from their mother cell (Figure 5C). We obtained similar results with *bud1*Δ mutant cells (Figure 5A-C). These results indicate that the ER diffusion barrier reinforces the confinement of the pheromone refractory state to the mother cell.

**Figure 5.**
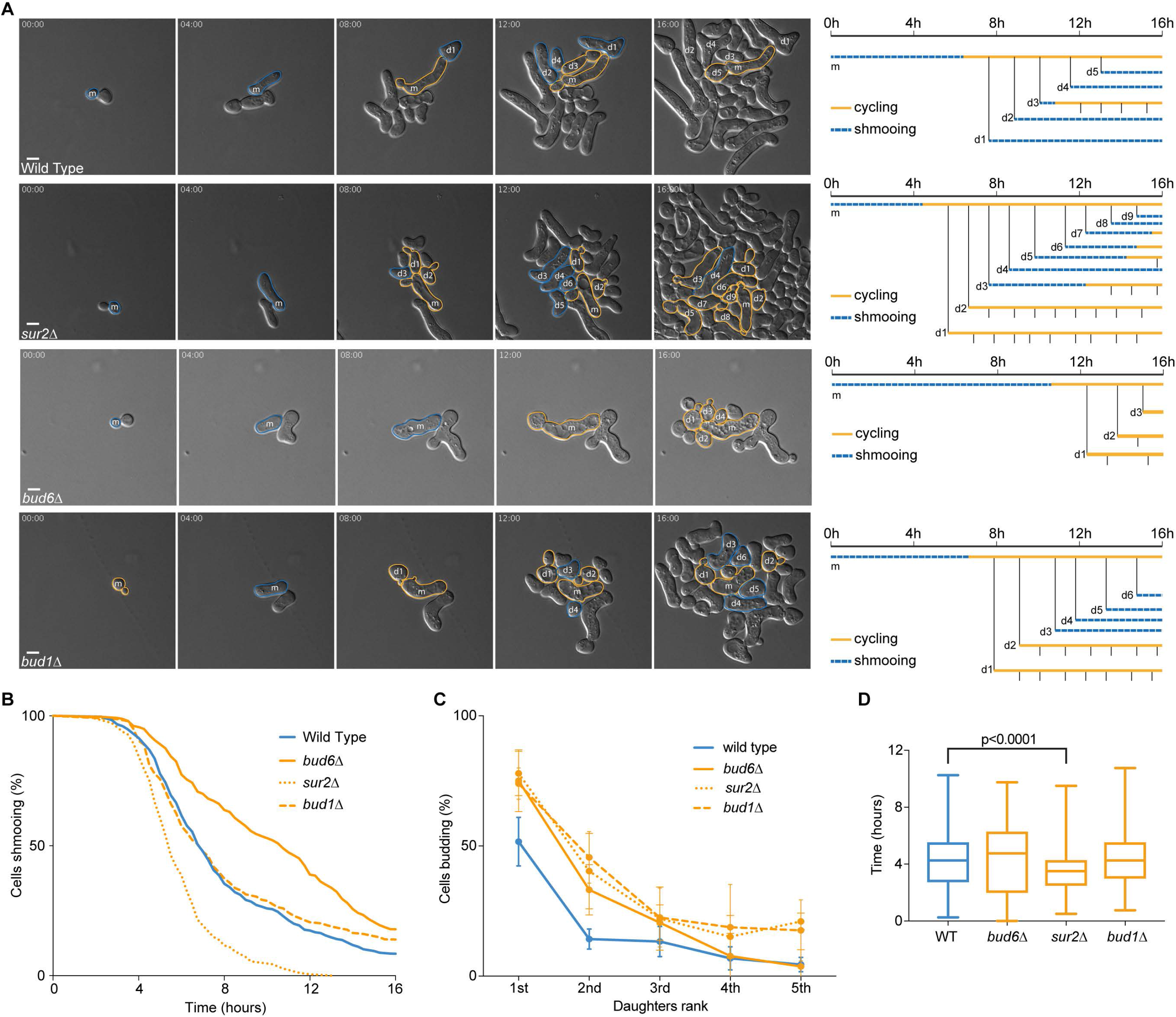
The ER diffusion barrier prevents daughter cells from inheriting the pheromone refractory state. (A) Escape of a wild-type, a *bud6*Δ, a *sur2*Δ and a *bud1*Δ cell exposed to 7 nM pheromone. (B) Percentage of initial cells still shmooing after the indicated time (n>154 cells). (C) Percentage of daughter cells budding immediately after separation from the mother cell (n>128 cells, n>121 cells, n>111 cells, n>88 cells and n>59 cells for the 1^st^, 2^nd^, 3^rd^, 4^th^ and 5^th^ daughter respectively). (D) Timing of escape of daughter cells that shmooed (*sur2*Δ is significantly different from wild type, p<0.0001, one-way ANOVA, all other comparisons are not significantly different, n=366 cells, 81 cells, 416 cells and 193 cells for wild-type, *bud6*Δ, *sur2*Δ and *bud1*Δ cells respectively).

Next, we wondered whether *bud6Δ, sur2*Δ or *bud1*Δ mutations had any effect on the timing of escape in the daughter cells that do shmoo in response to pheromone. We reasoned that these daughter cells may escape faster if they inherit factors promoting escape from their mother cell. However, we did not detect any general effect on the timing of daughter cells escape in these mutants (Figure 5D). We still observed that *sur2*Δ daughter cells escaped faster than wild type, *bud6*Δ and *bud1*Δ cells, as their mothers do.

Altogether, we conclude that diffusion barrier in the cortical ER helps confining the pheromone refractory state to the mother cell. Importantly, in cells lacking a functional barrier the inheritance of the refractory state by the daughter cells does not take place at the cost of the mother cell losing the pheromone refractory state. Indeed, *bud6*Δ, *sur2*Δ and *bud1*Δ mutant mother cells maintained this state as efficiently as wild type cells. Together these results indicate that the ER diffusion barrier confines the pheromone refractory state to the mother cell by preventing the formation of super-assemblies in daughter cells. Strikingly, this role of the barrier is most evident in the first few cycles after pheromone escape by the mother cell and much less so later on.

### The prion-like domain of Whi3 is required for symmetric inheritance of the refractory state in cells with an impaired diffusion barrier

We next reasoned that if inheritance of the pheromone refractory state by daughter cells depends on Whi3^mnem^ super-assemblies, then preventing the conversion of Whi3 to its mnemon form may lead to less daughter cells inheriting the pheromone refractory state upon diffusion barrier defects. To test this idea, we exposed *whi3-ΔpQ, whi3-ΔpQ bud6*Δ and *whi3-ΔpQ bud1*Δ mutant cells to pheromone and analyzed whether the daughters of cells escaping pheromone arrest were shmooing or budding after birth. Most *whi3*-Δ*pQ* daughter cells shmooed upon birth (Figure 6) and this was not significantly different in *whi3Δ-pQ bud6*Δ and *whi3Δ-pQ bud1*Δ cells. Taken together, these data argue that the Q-rich prion-like domain of Whi3 is involved in the inheritance of the pheromone refractory state by the daughters of cells that lack a diffusion barrier.

**Figure 6.**
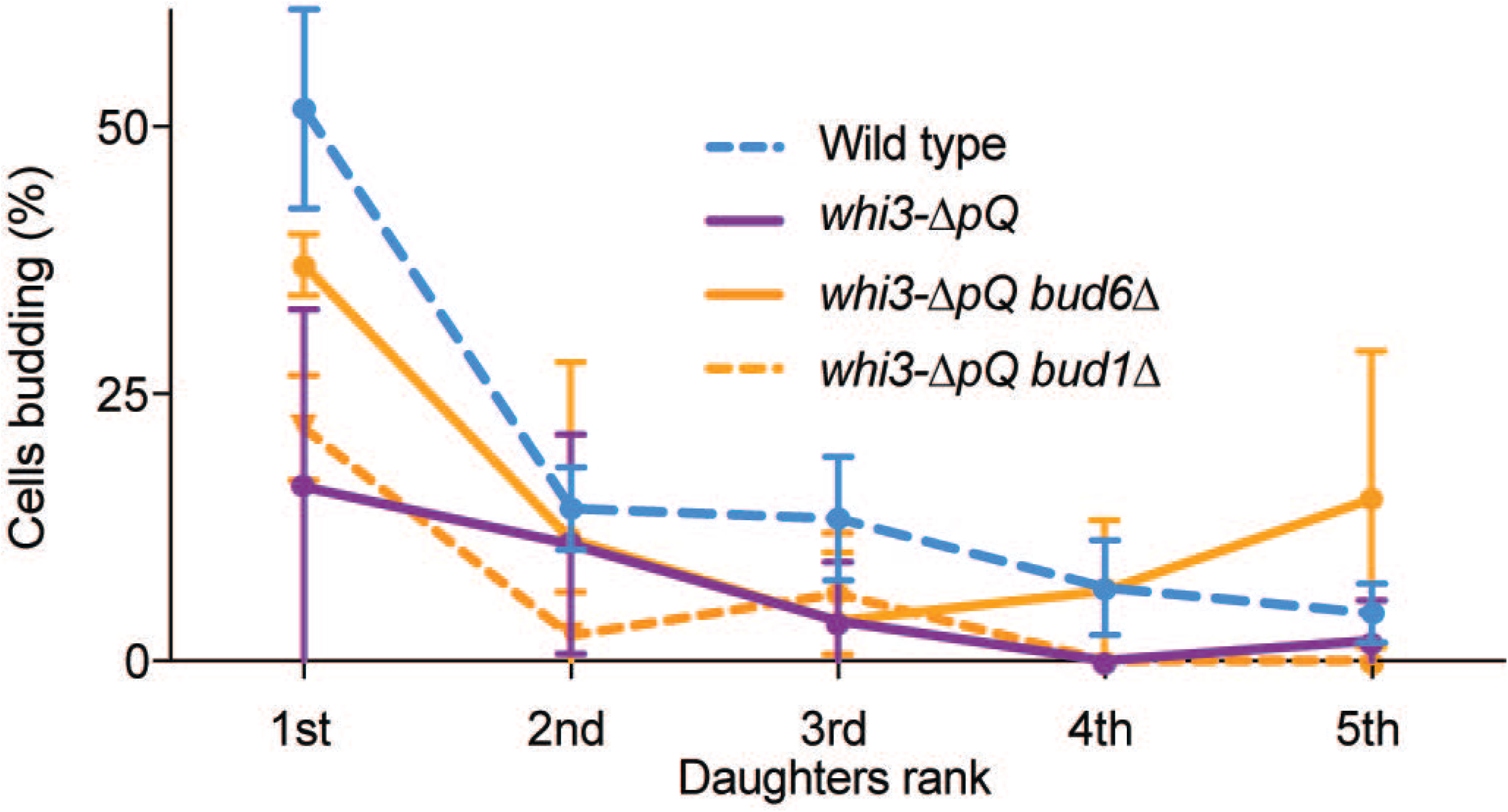
The polyQ domain of Whi3 is required for inheritance of the pheromone refractory state by diffusion barrier impaired daughter cells. Percentage of daughter cells of indicated genotypes budding immediately after separation from the mother cell. Mean ±SD are presented.

## Discussion

In this work, we investigated the mechanisms that allow the pheromone refractory state to be stable in the mother cells over many divisions while preventing it to being passed on to daughter cells. Our data support a model in which Whi3 adopts a self-templating conformation that depends on its prion-like domains upon prolonged pheromone exposure. The self-templating Whi3^mnem^ is confined by diffusion barriers to the mother cell and when this confinement is lost, Whi3^mnem^ does propagate to daughter cells. In a substantial fraction of the cases, it even starts to propagate in a mitotically stable manner, similarly to prions.

Indeed, we found that yeast cells can acquire a mitotically stable refractory state to pheromone and cells carrying this state were termed constitutive escapers. Some of these CE are small cell sized, phenocopying a loss of function of Whi3. In these small cell sized CE Whi3 forms foci, a hallmark of prion localization in yeast. Moreover, CE induction requires the prion-like domains of Whi3. These data indicate that Whi3 is in a dominant inactive, self-templating prion form in these CE variants. The fact that deleting *WHI3* increases CE frequency indicates that an important step of the transition to the CE state is the inactivation of Whi3 through its conversion to a prion form. Interestingly, the result that *whi3*Δ mutant cells are still able to acquire the CE state indicates the existence of additional prion-like factors in CE cells. Therefore, CE is a promising case for identifying novel prions in yeast.

During escape from pheromone arrest in diffusion barrier deficient cells, inheritance of the pheromone refractory state by daughter cells requires the prion like domains of Whi3, also indicating that Whi3^mnem^ is in a self-templating conformation. In diffusion barrier deficient cells, the pheromone refractory state is stable in the mother cells, suggesting that small assemblies are diffusing to daughter cells and seeding the pheromone refractory state there. However, in diffusion barrier defective cells only the first two to three daughter cells can inherit the pheromone refractory state efficiently. This is much less frequent in the next daughter cells. We propose that within the time of two to three cell cycles the super-assemblies mature to a more solid form that no longer generate seeds or captures them efficiently. These data provide experimental support for a mechanism of memory maintenance through a self-templating mechanism. This is a particularly interesting mechanism because it can establish a stable memory that lasts longer than the turnover of the Whi3 protein itself and prevents newly synthesized Whi3 to restore *WHI3* function.

All these data argue that Whi3 is able to form several types of assemblies, either being in a mnemon or in a prion form. Since, CE can have different cell sizes, it could also be that Whi3 can adopt several types of prion forms, which is reminiscent of the different strains prions can form (Derkatch et al., 1996). It may also be that the mnemon and the prion forms are actually the same and that what is lost are the mechanisms of confinement to the mother cell. CE were frequent in cells with defective diffusion barriers. Because *bud1*Δ cells display this feature, our results point to the ER diffusion barrier playing an important role in the prevention of CE induction. Indeed, only the ER diffusion barrier is affected by *BUD1* deletion, while the nuclear diffusion barrier remains intact (Clay et al., 2014). Yet, these observations beg the question of how diffusion barriers could prevent the induction of CE?

We found that [*PSI*+] prion induction and curing is not affected by whether the diffusion barriers are present or not. On the contrary, we observed that Whi3^mnem^ super-assemblies and the pheromone refractory state they encode are not confined as efficiently to the mother cell in diffusion barrier defective cells than in wild-type cells. Thus, we propose that one function of the diffusion barriers is to retain Whi3^mnem^ super-assemblies in the compartment where they are formed. Even though Whi3 is not known to be lipidated or having transmembrane domains, Whi3 associates with ER membranes (Vergés et al., 2007). Regardless of how it does so, such an association would allow the diffusion barrier to limit the diffusion of Whi3 seeds along ER membranes through the bud neck. More importantly, we found that Whi3^mnem^ super-assemblies are closely colocalizing with ER membranes during prolonged pheromone response. It will be therefore important in the future to understand how Whi3 is anchored at ER membranes to be able to test our model of retention by the ER diffusion barrier more thoroughly. Altogether a feature of mnemons may be their tight association with ER membranes allowing confinement to one cellular compartment. This is strikingly different to the Sup35 prion, most of which is detected in foci that are away from ER membranes. In this case, the presence of diffusion barriers in the ER membrane has no impact on exchange of seeds across the bud neck. Once Sup35 acquires its prion form in a cell, it can freely invade the bud and maintain the phenotypes associated with it in the progeny, and hence, the whole colony formed. We observed a similar localization pattern respective with ER membranes for Whi3 foci in CE cells. This suggests to us that the Whi3 prion form may differ from the mnemon form at least in part through their membrane association status. Whether the detachment of Whi3 in CE from ER membranes stems from a conformational change inhibiting Whi3 interaction with its ER anchor or from a change in the regulation of this interaction will need to be clarified.

Altogether, we propose that the absence of the barrier allows for the selection of stable prion strains of Whi3 and possibly other prions. The emergence of these prions would not be able to emerge if they are confined.

Many proteins can adopt prion-like behavior across all kingdom of life. Recent effort to characterize these proteins have suggested that this is probably more widespread than anticipated. We presented here that yeast cells have evolved mechanisms to confine and thus control some of them. We propose that this is not restricted to budding yeast. A case could be made for the protein CPEB, which adopt a prion-like conformation at activated dendritic spines (Khan et al., 2015; Si et al., 2010). Interestingly, dendritic spines are highly compartmentalized cellular appendages, and similarly to dividing yeast cells, their compartmentalization and morphology is controlled by septin proteins at the spine neck (Ewers et al., 2014). It may therefore be that CPEB prion conversion is confined to an activated dendritic spine through its compartmentalization by diffusion barriers, which could prevent the spreading of the prion form to the neighbouring non-activated spines. Another consideration may also be that confinement of prion-like elements such as mnemons may prevent their transformation into infectious prion particles. In this case, we could envision that pathologies involving prion-like behaviors may arise when confinement of these elements is lost.

## Material and Methods

### Strains

Strains used for escape from pheromone arrest were derivatives of the s288c BY4743 wild type (yYB4508: *MAT***a**, *his3Δ1 leu2Δ0, ura3Δ0 met15Δ0 lys2Δ0 ADE2 TRP1 bar1::kanMX*) with deletions obtained according to (Janke et al., 2004) (yYB4507: *sur2::NAT*; yYB4510: *bud6::NAT*; yYB4511: *bud1::NAT*). Strains used for Whi3 localization were derivatives of the wild type (yYB6520: *MAT***a***ura3Δ0, hisΔ13, leu2Δ0, TRP1, LYS2, ADE2, met15Δ0, WHI3-3GFP:kanMX, bar1::HIS3*) backcrossed in yYB4510 for *bud6*Δ (yYB10190), in yYB4507 for *sur2*Δ (yYB10192) or in yYB4511 for *bud1*Δ (yYB10187) strains. Strains for the co-localization of Whi3-3GFP and Sec61-mCherry were obtained by PCR tagging of Sec61 in yYB6520 (yFC203: *MAT***a***ura3Δ0, hisΔ13, leu2Δ0, TRP1, LYS2, ADE2, met15Δ0, WHI3-3GFP:kanMX,bar1::HIS3, Sec61-mCherry:kanMX)*. W303 strains used to test for [PSI+] induction and curing were obtained from Jonathan Weissman, yYB8040 (*MATα*, *leu2-3,-112; his3-11,-15; trp1-1; ura3-1; ade1-14; can1-100*, [*RNQ*+], [*PSI*+]). The *sur2*Δ strain was obtained by deleting *SUR2* according to Janke et al. (Janke et al., 2004) (yYB8435 *MATα*, *leu2-3,-112; his3-11,-15; trp1-1; ura3-1; ade1-14; can1-100, sur2::HIS3*, [*RNQ*+], [*PSI*+]). Localisation of Whi3 in *sur2*Δ^*CE*^ was analysed in strain yYB5147 (*his3Δ1, leu2Δ0, met15Δ0, ura3Δ0, LYS2, bar1::kanMX, sur2::NatMX, Whi3-GFP:HIS3MX6*). To analyse co-localisation of Sup35-GFP and Sec61-mCherry we used strain yFC202 (*MAT***a/alpha***SUP35/SUP35-GFP:HIS3; leu2-3,-112/leu2Δ0; his3-11,-15/his3Δ1; trp1-1/TRP1; ura3-1/ura3Δ0; ade1-4/ADE1; can1-100/CAN1, MET15/met15Δ0, SEC61/Sec61:mCherry:URA3* [*RNQ*+], [*PSI*+]). For stop codon readthrough, we used wild type (*MAT***alpha**, *leu2-3-112, his3-11,-15, trp1-1, ura3-1, ade1-4, can1-100*, [*RNQ*+], [*PSI*+], *pRS304 pGPD GST-UGA-GFP-pest:URA3*) or *sur2*Δ strain (*MAT***alpha**, *leu2-3-112, his3-11,-15, trp1-1, ura3-1, ade1-4, can1-100*, [*RNQ*+], [*PSI*+], *pRS304 pGPD GST-UGA-GFP-pest:URA3 sur2::KanMX*).

### Microscopy

All images were acquired either on a Personal Deltavision (Applied Precision) equipped with a CCD HQ2 camera (Roper) and 250W Xenon lamps controlled by Softworx or a Deltavision Elite (GE Healthcare) equipped with a sCMOS camera and solid-state light-emitting diodes controlled by Softworx. Fluorescein isothiocyanate and tetramethylrhodamine isothiocyanate filters were used for imaging GFP and mCherry fluorescence. Deconvolution was performed using Softworx.

### Microfluidics

Experiments were carried out with the ONIX microfluidic perfusion platform with Y04C microfluidic plates (CellAsic). Medium was yeast extract peptone dextrose (YPD) supplemented with 20 mg/ml casein and containing 7 nM α-factor (Figure 2 and 3E) or 6 nM α-factor (Figure 3C-D).

### Quantification of Shmooing

Shmooing or budding states were inspected visually. During microfluidic experiments, images were taken every 15 min. Unbudded cells that showed a polarized growth were counted as shmooing. Unbudded cells undergoing isotropic growth were counted as G1 cells. Usually these cells soon started forming a bud. Samples consisted of three independent clones.

### Quantification of Whi3-3GFP Super-Assemblies

Cells were grown in YPD supplemented with 20 mg/ml casein and containing 7 nM α-factor and were briefly centrifuged at 600 g, resuspended in SD-TRP medium, placed between slide and coverslip, and imaged immediately. Images were analyzed after deconvolution with Softworx software as before [27]. Three clones with total n > 122 cells were observed for each strain.

### Quantification of [PSI+] *de novo* appearance

[PSI+] [PIN+] wild type and *sur2*Δ cells were first cured with 3 passages on YPD agar medium containing 3 mM GuHCl. Red single colonies [psi-] [pin-] were assessed for their ability to become white again. Cells were plated on SC-Ade and YPD and the frequencies of appearance of white colonies were measured. White colonies were tested for their ability to become red again after passages on YPD containing 3 mM GuHCl.

### Quantification of [PSI+] curing during treatment with GuHCl

Cells were grown overnight in liquid YPD and diluted in the morning to OD_600nm_ = 0.2 in YPD with 3 mM GuHCl. Samples were taken every 30 minutes and plated on synthetic medium with low adenine concentration. Liquid cultures were kept in exponential phase during the experiment. Colonies were allowed to grow at 30°C for several days and the proportion of white and red colonies was assessed after 2 days of incubation at 4°C to allow for the red colour to develop well. We initially determined that curing started to happen after 12 hours of GuHCl treatment for both wild type and *sur2*Δ strains.

### Quantification of Stop-codon read-through by flow cytometry

For the stop codon read through experiments, wild-type [*PSI*+] and *sur2*Δ [*PSI*+] cells expressing chromosomally integrated pGPD GST-UGA-GFP-pest were grown for 5 hours without GuHCl or with 0.1mM or 1mM GuHCl. The GFP fluorescence intensity was measured with a BD Accuri C6 Flow Cytometer using 488 nm laser and 533/30 BD filter for 100 000 cells/ clone (3 clones each). The data was analyzed using FlowJo software (FlowJo LLC).

### Quantification of CE frequencies

Diploid strains heterozygous for the different mutations were sporulated (for example *SUR2/sur2*Δ or *WHI3/whi3*Δ). *MAT***a** spores were selected and their genotypes determined by growth on selection media. Strains were grown in YPD until mid-log phase and spotted on solid YPD and solid YPD containing 0.6μM α-factor. Colonies were counted after 2-3 days of growth at 30°C.

The analysis was conducted on the estimates of the of relative yeast density obtained from the colony count (C) at an appropriate dilution (D). The relative density for each clone was obtained in the presence of the pheromone (p) and the corresponding control (c), so the relative performance is given by the ratio:

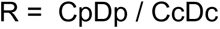

The ratio was logit transformed in an attempt to standardize the variance. Having fitted the average for each yeast strain, the residuals were clearly strongly asymmetrical (Supplemental Figure 1 shows the deviation from a cumulative normal distribution). This pattern might be expected if in a subset of cases the constitutive escaper phenotype occurred early in the culture. The departure from a cumulative normal distribution of residuals is abrupt for standardized residuals greater than one (shown by the vertical line). Since the incidence of these outcomes did not differ between strains (χ^2^ = 3.02, P = 0.88) they were excluded from subsequent analysis of the ratios.

Figure 1C shows the distribution of the logit transformed ratios (R) for each genotype. A linear model describing the means of each strain, showed a highly significant difference between the three strains for which there were a priori expectations of a stronger effect of the pheromone (*whi3*-Δ*pQ*, *bud1Δ whi3-ΔpQ* and the wildtype) and the remainder (ANOVA P < 2e-16). The fitted values for these two categories is shown by the red line. There were no significant differences between the means for this remaining group (P = 0.41) whereas there were significant differences among the three (P < 0.0007) - in particular *whi3*-Δ*pQ* was markedly lower than the wild type (P<0.0008). The R package for this analysis is available in the supplementary material.

### Cell sizes measurements

Cell sizes were determined using a CASY cell counter model TTC (Schärfe system). Strains were grown to early log phase, diluted in CASYton (Electrolyte buffer from Schärfe system) and processed according to the manufacturer instructions.

### Quantification of Sup35 foci and Whi3 super-assemblies/foci/granules co-localisation with ER membranes

For Whi3-3GFP localisation, cells were grown in YPD supplemented with 20 mg/ml casein and containing 7nM α-factor and were briefly centrifuged at 600 g, resuspended in SD-TRP medium, placed on a SC-TRP agar pad covered by a coverslip, and imaged immediately. Images were analysed after deconvolution with Softworx software. Three clones with total n = 194 cells, 250 cells, 170 cells and 153 cells for 3 hours, 4 hours, 6 hours and 10.5 hours condition were observed for each strain. A total of 333, 402, 393 and 317 super-assemblies were analysed at 3 hours, 4 hours, 6 hours and 10.5 hours’ time points. Note that we only analysed super-assemblies that were in the 5 best focal planes as co-localisation was difficult to assess on the top and bottom focal planes and we only counted Sup35-GFP foci and Whi3-3GFP super-assemblies in the mother cells.

## Acknowledgements

We would like to thank Robin Hannay for his help in creating some of the strains. This work was supported by a Biotechnology and Biological Sciences Research Council Project grant (BB/S001204/1) to FC, QMUL to FC and YL, the Academy of Finland (317038) and Sigrid Jusélius Foundation to JS and the European Research Council (BarrAge 250278) and ETH Zurich (to J.S., F.C. and Y.B.).

---

**Supplemental Figure 1.**
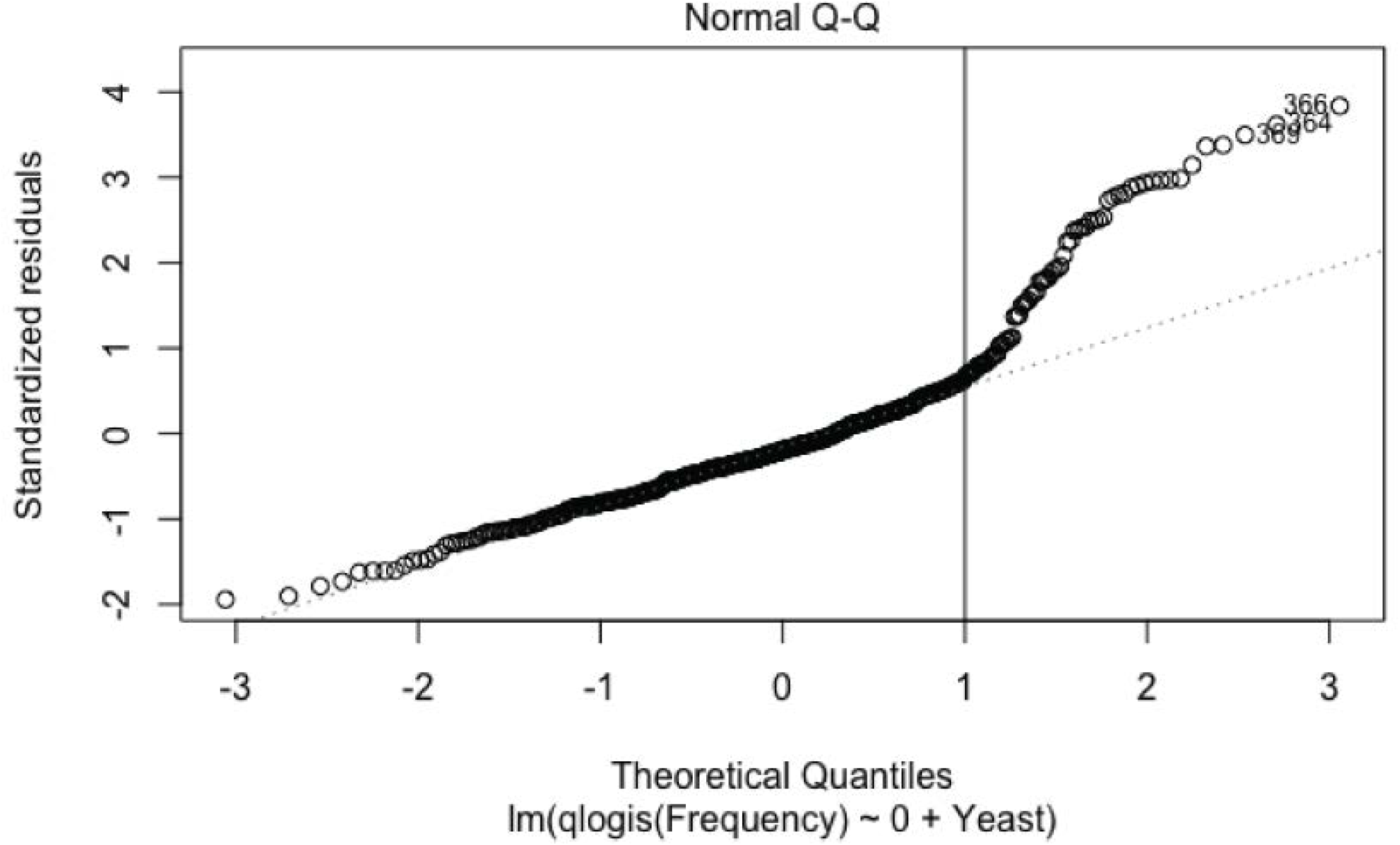
Deviation of the residuals from a cumulative normal distribution.

**Supplemental Figure 2.**
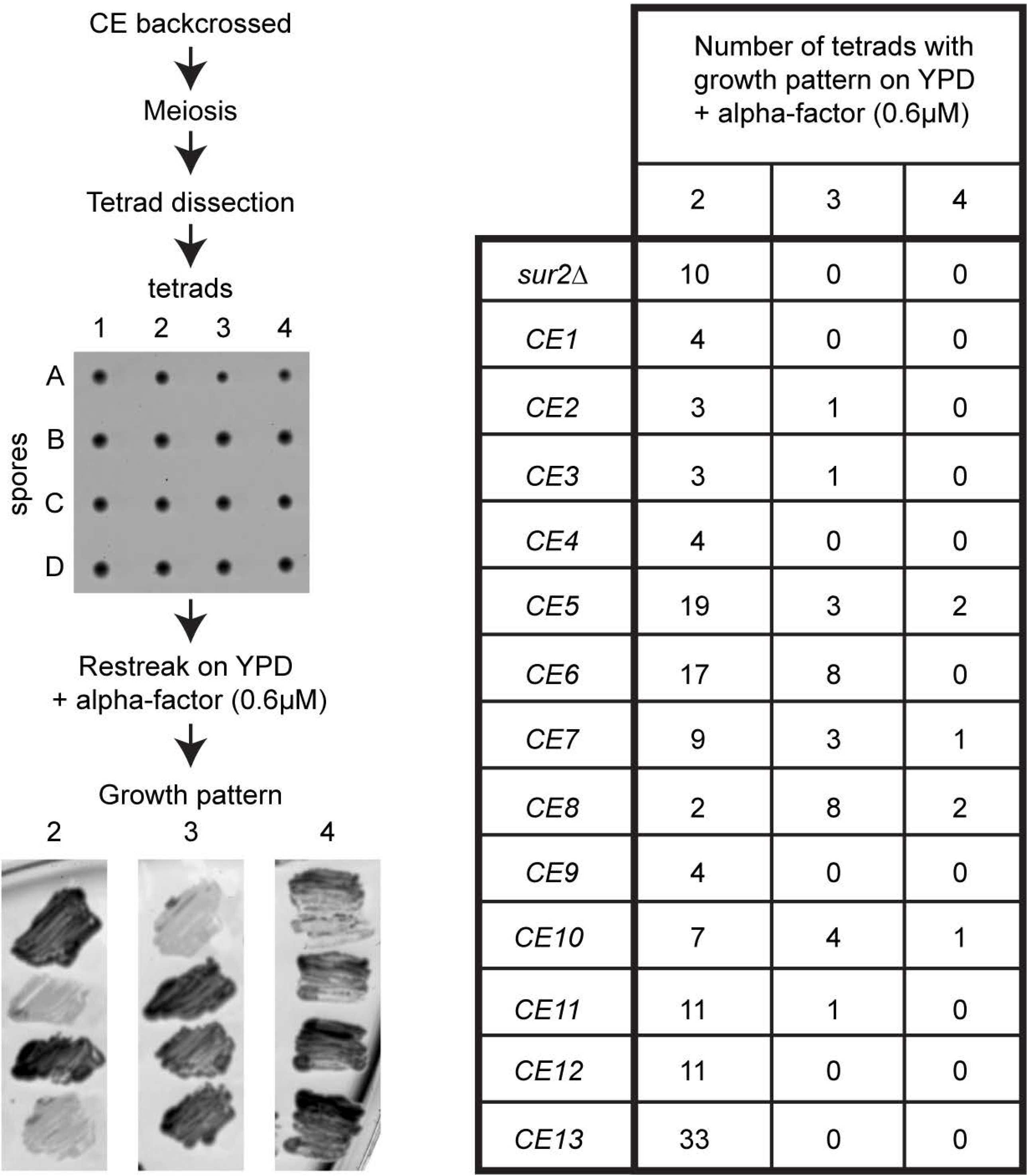
Non-mendelian inheritance of the CE phenotype during meiosis. Schematic of the experimental test (left) and results (right) of 2 backcrosses of *sur2*Δ parental strain (pooled in one line) and 13 CE backcrossed.

**Supplemental Figure 3.**
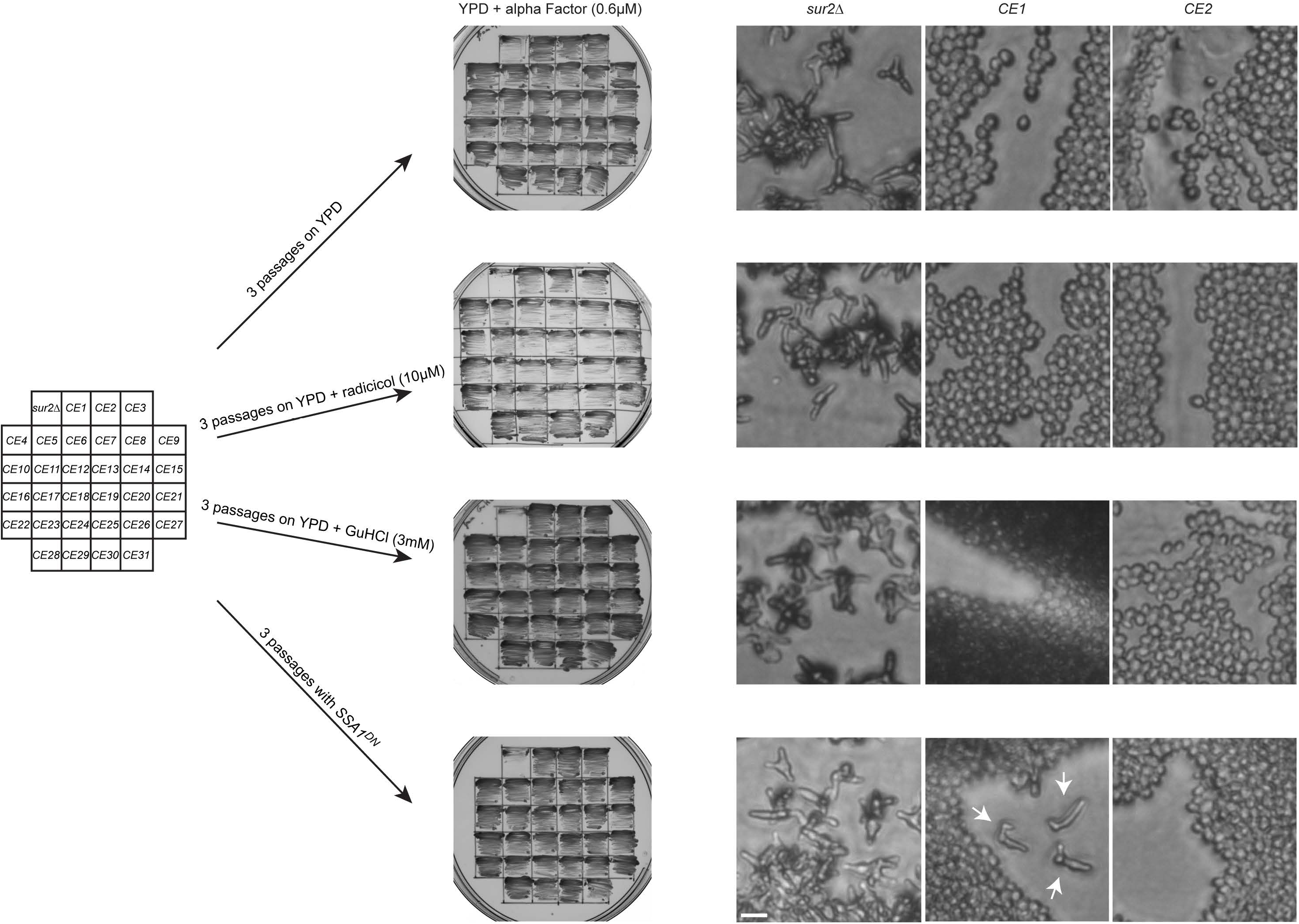
The CE phenotype can be partially cured by Ssa1 inhibition. Schematic of the experimental test. CE were isolated, restreaked 3 times on YPD, YPD + radicicol (10μM) or YPD + GuHCl (3mM) or transformed with a plasmid expressing *SSA1*^*DN*^ and restreaked 3 times on selective medium. CE were then tested again on YPD + alpha factor (0.6μM). The right panels display microscopic images of the parental *sur2*Δ strain and CE1 and CE2 on YPD + alpha factor (0.6μM) after the corresponding treatments. Arrows point at shmooing cells in CE1 transformed with a plasmid expressing *SSA1^DN^*. Scale bar = 10μm.

